## Supplementary material for "A BAC-guided haplotype assembly pipeline increases the resolution of the virus resistance locus *CMD2* in cassava": Supp file 1

### Supplemental Figures and Tables

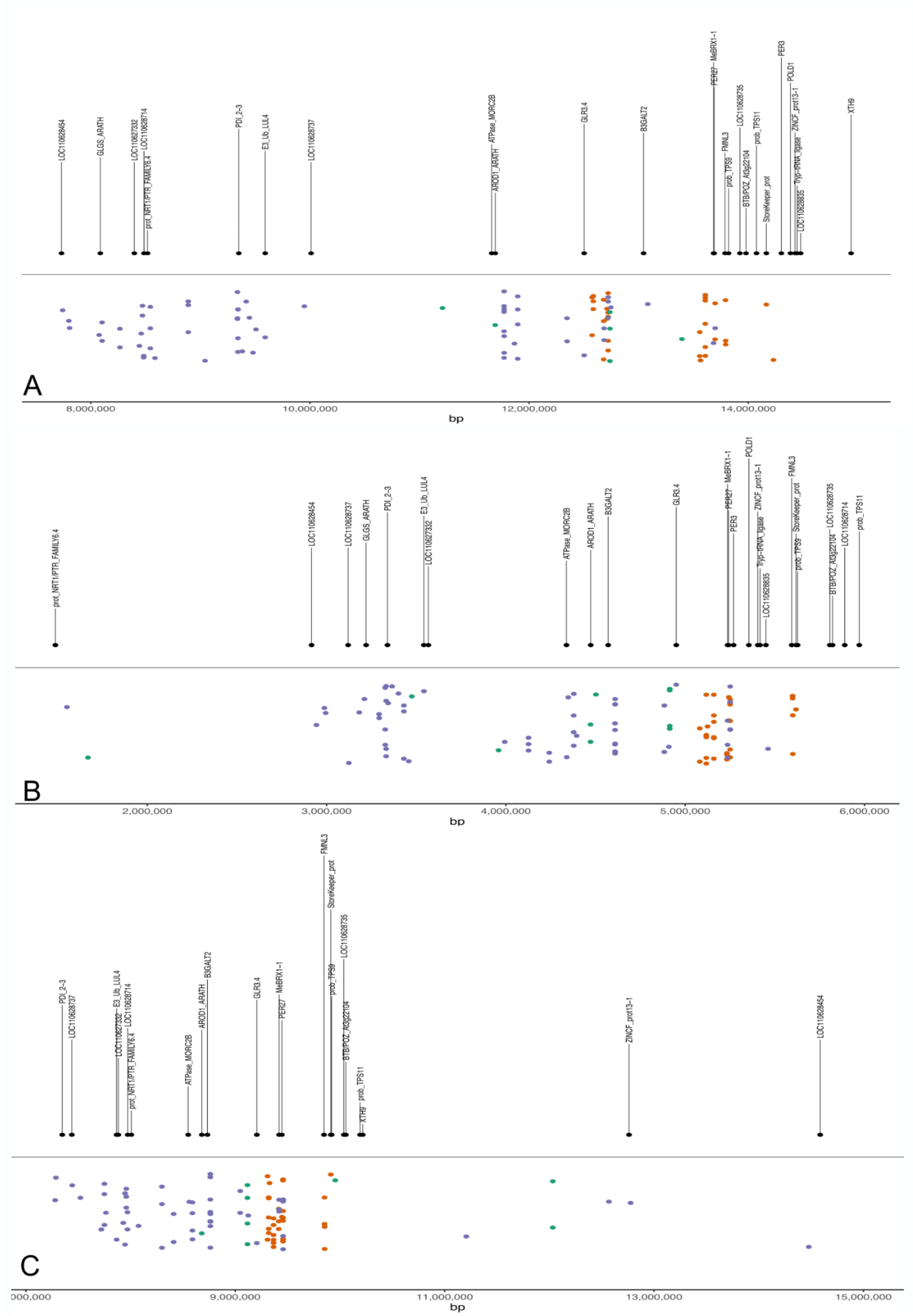

**Supplemental Figure 1: Markers and genes mapping on 60444 CMD2.** The *CMD2* locus in 60444 GCA_003957885.1 Kuon (A), 60444 GCA_963409065.1 Cornet haplotype 4632 (B), and 60444 GCA_963409065.1 Cornet haplotype 100051 (C). The black dots (above the horizontal black line) show the alignment positions of genes that can be found in the *CMD2* region. The colored dots (below the horizontal black line) indicate various molecular markers associated with CMD resistance. The green dots indicate classical markers (RFLP and SSR markers) published by Akano et al. 2002 (43), Lokko et al. 2005 (44), Okogbenin et al. 2007 (45) and Okogbenin et al. 2012 (46). Orange dots indicate CMD2 SNP markers published by Rabbi et al. 2022 (12), and violet dots indicate markers published by Wolfe et al. 2016 (11). The x-axis of the plots indicate the base pair (bp) position on the genome.

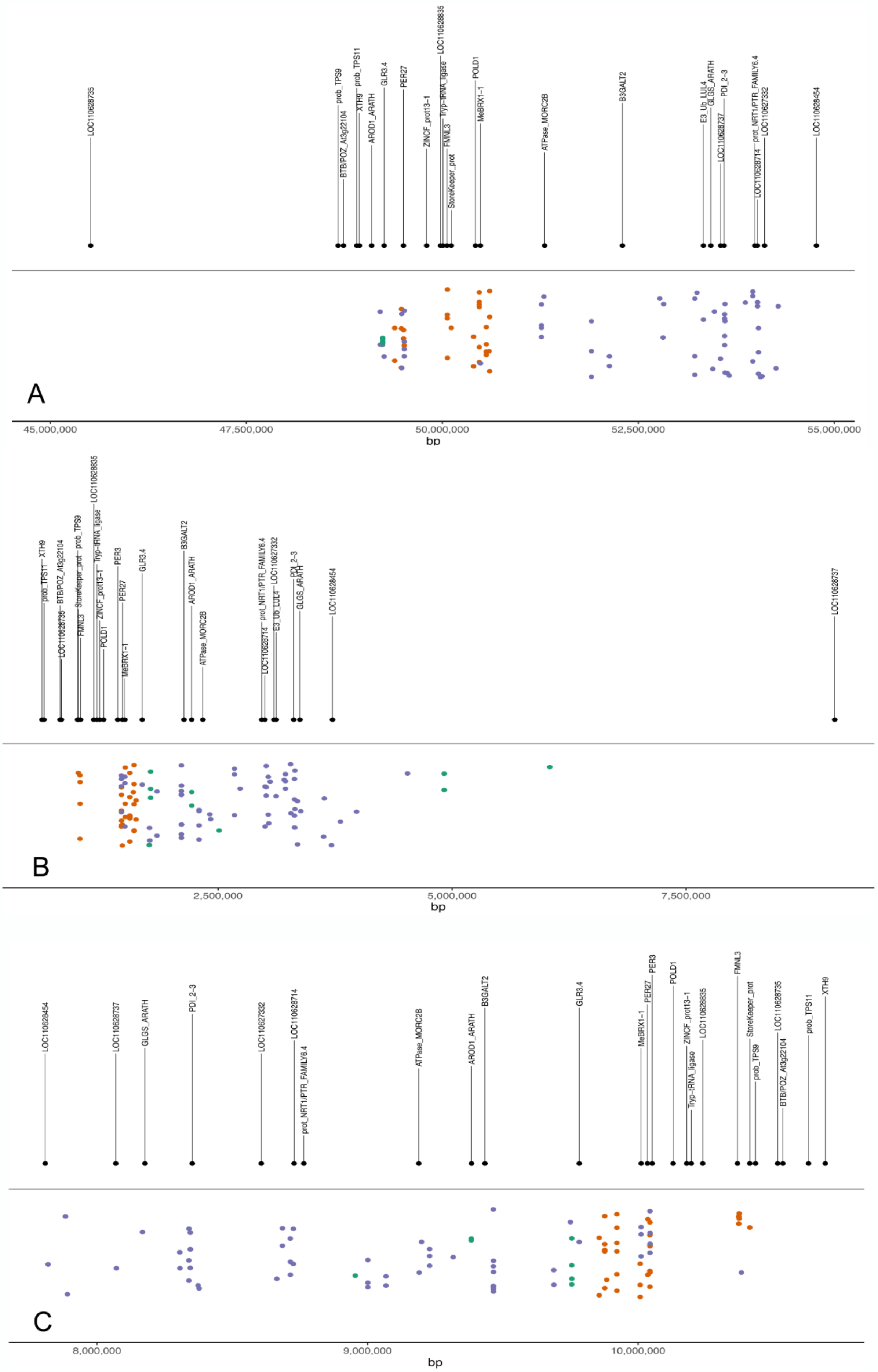

A
LOC110628735
BTB/POZ_At3g22104
prob_TPS9
XTH9
prob_TPS11
AROD1_ARATH
GLR3.4
PER27
ZINCF_prot13-1
LOC110628835
Tryp-tRNA_ligase
FMNL3
StoreKeeper_prot
POLD1
MeBRX1-1
ATPase_MORC2B
B3GALT2
E3_Ub_LUL4
GLGS_ARATH
LOC110628737
PDI_2-3
LOC110628714
prot_NRT1/PTR_FAMILY6.4
LOC110627332
LOC110628454
45,000,000
47,500,000
50,000,000
52,500,000
55,000,000
bp
B
prob_TPS11
XTH9
LOC110628735
BTB/POZ_At3g22104
FMNL3
StoreKeeper_prot
prob_TPS9
ZINCF_prot13-1
Tryp-tRNA_ligase
LOC110628835
POLD1
PER3
PER27
MeBRX1-1
GLR3.4
B3GALT2
AROD1_ARATH
ATPase_MORC2B
LOC110628714
prot_NRT1/PTR_FAMILY6.4
E3_Ub_LUL4
LOC110627332
PDI_2-3
GLGS_ARATH
LOC110628454
LOC110628737
2,500,000
5,000,000
7,500,000
bp
C
LOC110628454
LOC110628737
GLGS_ARATH
PDI_2-3
LOC110627332
LOC110628714
prot_NRT1/PTR_FAMILY6.4
ATPase_MORC2B
AROD1_ARATH
B3GALT2
GLR3.4
MeBRX1-1
PER27
PER3
POLD1
ZINCF_prot13-1
Tryp-tRNA_ligase
LOC110628835
FMNL3
StoreKeeper_prot
prob_TPS9
LOC110628735
BTB/POZ_At3g22104
prob_TPS11
XTH9
8,000,000
9,000,000
10,000,000
bp

**Supplemental Figure 2: Markers and genes mapping on *CMD2* from TME3 genome.** The CMD2 locus in TME3 GCA_003957885.1 Kuon (A), TME3 GCA_963409055.1 Cornet haplotype 100036 (B), and GCA_963409055.1 Cornet haplotype 100004 (C). The black dots (above the horizontal black line) show the alignment positions of genes that can be found in the CMD2 region. The colored dots (below the horizontal black line) indicate various molecular markers associated with CMD resistance. The Green dots indicate classical markers (RFLP and SSR markers) published by Akano et al. 2002 (43), Lokko et al. 2005 (44), Okogbenin et al. 2007 (45) and Okogbenin et al. 2012 (46). Orange dots indicate CMD2 SNP markers published by Rabbi et al. 2022 (12), and violet dots indicate markers published by Wolfe et al. 2016 (11). The x-axis of the plots indicates the base pair (bp) position on the genome.

| **Probe** | **Primer** | **Sequence** |
|---|---|---|
| PDI | PDI_F | ACAAGATGGACAAACGTCCT |
| | PDI_R | TGAATGTGAAGAAAGGAGCA |
| Snp1 | snp1_F | TGAGCCATGAATGATGTTGTG |
| | snp1_R | TGGTTGATGGAAGATCAGCA |
| Snp3 | snp3_F | AAATCTTGTTCGCGTCCTTC |
| | snp3_R | GGTATCCAACTCCCCGAAAT |
| 169 | 169_F | ATCATCACCATGAGCAACGA |
| | 169_R | CAATGGTCCTCAGGAATGCT |
| 158 | 158_F | GAGATCCACGATGATGCTGA |
| | 158_R | TCAAAGCAACAAGCAAGGAA |
| PDI3600 | pdi3600_F | ATCACTAGCAGGGGCAAAAA |
| | pdi3600_R | ACTTGCCAAATTTGATCTTCA |
| PDI3100 | pdi3100_F | TGTTACCAGGAAAAGAGTTCTGC |
| | pdi3100_R | GCGAGCTGCAGTTGCTAAAT |

| | | |
|---|---|---|
| 3 | 3_F | GTGGTTATTTGGCAACATGG |
| | 3_R | TTGAATTTAGCATTCCGTTGAA |
| 32 | 32_F | GACAGCTGCTATACCTATCTTGG |
| | 32_R | TAACTTTACCAATAAAAGCCAATCC |
| 80 | 80_F | TTATCTGGATATGGGGTGGA |
| | 80_R | AAGGAAAAGGGGCAGGATTA |
| Z3 | Z3F | AAAGTTTGAGTTTCGGCTCTGTCC |
| | Z3R | GAAAGACTATCTCACAACCGGG |
| Z7 | Z7F | TAGGGATTATGTCTTGGCTGTC |
| | Z7R | CTTTAGTGTCAATTCCCTGTGG |
| Z9 | Z9F | CAACTCCTTTCCACTCCTCAATG |
| | Z9R | CAAACCCAACAAGTCCAGCAGATT |
| Z12 | Z12F | CAGACAGCTTGTCCCTACTAAATC |
| | Z12R | CCAGGAATTTCACTAGAAGAAGGG |
| Z16 | Z16F | CAACAGGAATCAAGTCACTCAC |
| | Z16R | GGTTCTTCCTTCAACTCCTCTGAT |
| Z18 | Z18F | AGAGGTCTTCAGTTGGTTATGG |
| | Z18R | CAAATCCAACCCAGGAACCTCTT |
| Z21 | Z21F | TTTGGGATACAGGGTTGCTTAG |
| | Z21R | GATTTGTGAAGACGAAGGCTCT |
| Z22 | Z22F | AGCTCATGTGAGGGAGTTGGATAG |
| | Z22R | CATCCTCCTCTCTGATTCTCAA |
| Z23 | Z23F | CACCTCCACCATACTATCCAAATC |
| | Z23R | GCAGCAACTCCATAAACCTGTCA |

| | | |
|---|---|---|
| Z30 | Z30F | TTTCCATCTTGACTCTGCCTCC |
| | Z30R | CCATCTCAAGAAAGGTTGCTAC |
| Z36 | Z36F | ACCTTCAAGAACCAAGCTGATG |
| | Z36R | GAGATGCAGGATATGTGCCTGA |
| Z37 | Z37F | AATAGACCTGTGCATCAGGGTG |
| | Z37R | CTTGCTGCTTACCGAGTAGATGA |
| Z39 | Z39F | TGCATCTCAATCTCCTGGTATC |
| | Z39R | GAATCGAGTCTCACCTACAAGAG |
| Z41 | Z41F | TAGCAGTTGCTGTGGGTTACAATG |
| | Z41R | GAATGTCTGAAATGGGCTACTTGG |
| Z86 | Z86F | GGCTAATCCAGCAAGAAAGAGAACG |
| | Z86R | CACCCGTAGAAAGAAAGACCCGAAT |
| Z87 | Z87F | CAACTGGAAGGAAGGATGACAGAG |
| | Z87R | CAGCATTGATTTGTCCTTGTCC |
| Z91 | Z91F | TTATAGAGGTCAGGGTGGTTCTTGC |
| | Z91R | CAGCCGCGTGAATCAACAGTAAAG |

#### Supplemental Table 1: Primers used in this study for making BAC probes

| Genomes | BAC mapping percentage | BUSCO completeness % |
|---|---|---|
| **60444 with reads above 8kb, Flye default setting** | **81.45** | **72,9** |
| GCA_003957885.1 (60444) | 73.15 | 98,6 |
| 60444 all reads, Flye default setting | 81.40 | 70,2 |
| 60444 all reads,-L3000, wtdbg2 default setting | 53.09 | 77,5 |
| 60444 all reads,L10000, wtdbg2 default setting | 56.55 | 75,9 |
| 60444 all reads,L2000, wtdbg2 default setting | 59.81 | 11,3 |
| 60444 all reads,L5000, wtdbg2 default setting | 63.30 | 76,8 |
| 60444 all reads,-L8000, wtdbg2 default setting | 44.90 | 76,2 |
| 60444 all reads,L300, wtdbg2 default setting | 46.50 | 70,1 |

| | | |
|---|---|---|
| 60444 all reads, CANU with stopOnLowCoverage=5 and cnsErrorRate=0.25 | 67.46 | 98 |
| 60444 with reads above 8kb, CANU with stopOnLowCoverage=5 and cnsErrorRate=0.25 | 62.34 | 88,3 |
| **TME3 with reads above 8kb, Flye default setting** | **80.02** | **95,8** |
| GCA_003957995.1 (TME3) | 61.08 | 98,2 |

#### Supplemental Table 2: BAC mapping percentage and BUSCO completeness on genome assemblies.

| **Genomes** | **BUSCO** | | **QUAST** | | | |
|---|---|---|---|---|---|---|
| | **Single** | **Duplication** | **Total length (Gb)** | **N50** | **#Contigs** | **N percentage** |
| cv. 60444 | 57 % | 15.9 % | 1,42 | 18240505 | 6402 | 16039.45 |
| TME 3 | 60.6 % | 35.2 % | 1,56 | 22737009 | 8161 | 16936.51 |
| GCA_003957885.1 (60444) | 61,2 % | 37 % | 1,27 | 59195861 | 4440 | 23496.76 |
| GCA_003957995.1 (TME3) | 55.2 % | 43.4 % | 1,22 | 53513187 | 5398 | 22496.91 |

#### Supplemental Table 3: Statistics of genome assemblies.

| **Gene_ID** | **prped_probability** | **60444_presence (GCA_9634090 65.1 Cornet)** | **TME3-NCBI_presence (GCA_0039579 95.1 Kuon)** | **60444_Isoseq** | **TME3_Isoseq** | **CMD2_candidate** | **Length** | **Interproscan_domain** |
|---|---|---|---|---|---|---|---|---|
| evm.model.Hap2-Super-Scaffold_100004.552 | 0.9010995973506256 | Out CMD2 contig | V | V | X | X | 868 | GluR_Plant<br>Periplasmic binding protein-like II<br>Receptor family ligand binding region<br>Bacterial extracellular solute-binding proteins- family 3<br>GLUTAMATE RECEPTOR 3.4<br>GluR_14<br>Ligand-gated ion channel<br>PBP1_GABAb_receptor_plant<br>IGluLR_plant<br>Periplasmic binding protein-like I<br>IONOTROPIC GLUTAMATE RECEPTOR |

| | | | | | | | |
|---|---|---|---|---|---|---|---|
| evm.model.Hap1-Super-Scaffold_100036.36 | 0.8557790799817562 | V | X | X | V | V | 896 | Ligand-gated ion channel<br>Periplasmic binding protein-like II<br>PBP1_GABAb_receptor_plant<br>Metabotropic gamma-aminobutyric acid type B receptor signature<br>IGluLR_plant<br>IONOTROPIC GLUTAMATE RECEPTOR<br>Receptor family ligand binding region<br>GLUTAMATE RECEPTOR 3.4<br>Periplasmic binding protein-like I<br>Bacterial extracellular solute-binding proteins- family 3<br>GluR_14<br>consensus disorder prediction |
| evm.model.Hap2-Super-Scaffold_100004.551 | 0.8395016558903129 | V | V | V | V | X | 911 | Bacterial extracellular solute-binding proteins- family 3<br>Receptor family ligand binding region<br>IGluLR_plant<br>GluR_Plant<br>IONOTROPIC GLUTAMATE RECEPTOR<br>GLUTAMATE RECEPTOR 3.7<br>PBP1_GABAb_receptor_plant<br>Prokaryotic membrane lipoprotein lipid attachment site profile.<br>Periplasmic binding protein-like I<br>Ligand-gated ion channel<br>Metabotropic gamma-aminobutyric acid type B receptor signature<br>Periplasmic binding protein-like II<br>Metabotropic glutamate GPCR signature |
| evm.model.Hap2-Super-Scaffold_100004.501 | 0.8114243528832168 | Out CMD2 contig | Out CMD2 contig | V | V | X | 821 | consensus disorder prediction<br>NODULIN HOMEOBOX |
| evm.model.Hap2-Super-Scaffold_100004.497 | 0.7893735297342348 | V | V | X | X | V | 589 | MEMBRANE PROTEIN |
| evm.model.Hap1-Super-Scaffold_100036.99 | 0.7733181488118763 | V | V | V | V | X | 925 | CBM20 (carbohydrate binding type-20) domain profile.<br>Ribonuclease E/G family<br>CBM_20_2<br>RIBONUCLEASE E/G-LIKE PROTEIN-CHLOROPLASTIC<br>RNaseEG: ribonuclease- Rne/Rng family<br>Nucleic acid-binding proteins<br>Immunoglobulins<br>Starch-binding domain-like |
| evm.model.Hap2-Super-Scaffold_100004.541 | 0.7270525556199235 | X | X | V | X | X | 729 | 35exoneu6<br>POLYMYOSITIS/SCLERODERMA AUTOANTIGEN-RELATED<br>Ribonuclease H-like<br>HRDC domain<br>HRDC-like<br>PROTEIN RRP6-LIKE 3<br>3'-5' exonuclease |

| | | | | | | | | |
|---|---|---|---|---|---|---|---|---|
| evm.model.Hap2-Super-Scaffold_100004.545 | 0.7208088129583924 | V | V | X | X | V | 485 | Terpene synthase family- metal binding domain<br>Terpene Cyclase Like 1 C Terminal Domain<br>Farnesyl Diphosphate Synthase<br>Isoprenoid Synthase Type I<br>TERPENE SYNTHASE 12-RELATED<br>Terpene synthase- N-terminal domain<br>Terpenoid synthases<br>Terpenoid cyclases/Protein prenyltransferases<br>OS04G0344100 PROTEIN-RELATED |
| evm.model.Hap1-Super-Scaffold_100036.47 | 0.7152823677851056 | V | V | X | X | V | 408 | Terpene_cyclase_plant_C1<br>Terpene Cyclase Like 1 C Terminal Domain<br>OS04G0344100 PROTEIN-RELATED<br>Terpenoid cyclases/Protein prenyltransferases<br>Farnesyl Diphosphate Synthase<br>Isoprenoid Synthase Type I<br>Terpene synthase family- metal binding domain |
| evm.model.Hap2-Super-Scaffold_100004.525 | 0.7071564923891883 | X | X | V | V | X | 765 | Late exocytosis- associated with Golgi transport<br>Calcium-dependent channel- 7TM region-putative phosphate<br>consensus disorder prediction<br>Cytosolic domain of 10TM putative phosphate transporter<br>PROBABLE MEMBRANE PROTEIN DUF221-RELATED<br>consensus disorder prediction<br>PROTEIN OSCA1 |
| evm.model.Hap1-Super-Scaffold_100036.68 | 0.6951134205709454 | V | V | X | X | V | 754 | consensus disorder prediction<br>Late exocytosis- associated with Golgi transport<br>Calcium-dependent channel- 7TM region-putative phosphate<br>PROTEIN OSCA1<br>PROBABLE MEMBRANE PROTEIN DUF221-RELATED<br>Cytosolic domain of 10TM putative phosphate transporter |
| evm.model.Hap1-Super-Scaffold_100036.73 | 0.6541367694570865 | V | X | X | V | V | 562 | POT family<br>MFS general substrate transporter like domains<br>PROTEIN NRT1/ PTR FAMILY 6.4-LIKE<br>OLIGOPEPTIDE TRANSPORTER-RELATED |
| evm.model.Hap2-Super-Scaffold_100004.537 | 0.6116018128562075 | X | X | V | V | X | 366 | Prephenate dehydratase domain profile.<br>ACT_CM-PDT<br>Prephenate dehydratase signature 2.<br>PBP2_Ct-PDT_like<br>Periplasmic binding protein-like II<br>AROGENATE/PREPHENATE DEHYDRATASE<br>ACT domain profile.<br>ACT-like<br>Prephenate dehydratase<br>PREPHENATE DEHYDRATASE P PROTEIN |

| | | | | | | | | |
|---|---|---|---|---|---|---|---|---|
| evm.model.Hap2-Super-Scaffold_100004.521 | 0.5979740774809379 | Out CMD2 contig | V | V | V | X | 589 | MFS general substrate transporter like domains<br>PROTEIN NRT1/ PTR FAMILY 6.4-LIKE<br>OLIGOPEPTIDE TRANSPORTER-RELATED<br>POT family<br>MFS general substrate transporter |
| evm.model.Hap1-Super-Scaffold_100036.62 | 0.5967214267801142 | Out CMD2 contig | X | V | V | X | 686 | CW-type Zinc Finger<br>Histidine kinase-- DNA gyrase B-- and HSP90-like ATPase<br>ZINC FINGER CW-TYPE COILED-COIL DOMAIN PROTEIN 3.<br>OS06G0622000 PROTEIN<br>consensus disorder prediction<br>ATPase domain of HSP90 chaperone/DNA topoisomerase II/histidine kinase<br>Morc6 ribosomal protein S5 domain 2-like |
| evm.model.Hap1-Super-Scaffold_100036.55 | 0.5716795773591182 | X | X | X | X | V | 879 | SNF2 family N-terminal domain<br>TRANSCRIPTION TERMINATION FACTOR 2-RELATED<br>RING/U-box<br>Zinc/RING finger domain<br>RING-type zinc-finger<br>Superfamilies 1 and 2 helicase ATP-binding type-1 domain profile.<br>Superfamilies 1 and 2 helicase C-terminal domain profile.<br>SF2_C_SNF<br>consensus disorder prediction<br>DEXDc_SHPRH-like<br>helicmild6<br>P-loop containing nucleoside triphosphate hydrolases<br>ultradead3<br>Helicase conserved C-terminal domain<br>ring_2 |
| evm.model.Hap2-Super-Scaffold_100004.507 | 0.5694020177734762 | V | V | V | V | X | 437 | cax: calcium/proton exchanger<br>VACUOLAR CALCIUM ION TRANSPORTER<br>Sodium/calcium exchanger protein<br>caca2: calcium/proton exchanger |
| evm.model.Hap2-Super-Scaffold_100004.539 | 0.5426593147869594 | V | V | X | X | V | 351 | PROSTAGLANDIN REDUCTASE<br>N-terminal domain of oxidoreductase<br>NAD(P)-binding Rossmann-fold domains<br>2-ALKENAL REDUCTASE (NADP(+)-DEPENDENT)-LIKE<br>Zinc-binding dehydrogenase<br>GroES-like<br>PKS_ER_names_mod |

| | | | | | | | | |
|---|---|---|---|---|---|---|---|---|
| evm.model.Hap2-Super-Scaffold_100004.542 | 0.5280101953343767 | X | X | X | X | V | 529 | Farnesyl Diphosphate Synthase<br>Isoprenoid Synthase Type I<br>TERPENE SYNTHASE 12-RELATED<br>Terpene synthase family- metal binding domain<br>Terpene_cyclase_plant_C1<br>Terpenoid synthases<br>Farnesyl Diphosphate Synthase<br>Terpene synthase- N-terminal domain<br>OS04G0344100 PROTEIN-RELATED<br>Terpene Cyclase Like 1 C Terminal Domain<br>Terpenoid cyclases/Protein prenyltransferases |
| evm.model.Hap2-Super-Scaffold_100004.532 | 0.5277047250642454 | V | V | X | V | V | 613 | Morc6 ribosomal protein S5 domain 2-like<br>ZINC FINGER CW-TYPE COILED-COIL DOMAIN PROTEIN 3.<br>Histidine kinase-- DNA gyrase B-- and HSP90-like ATPase<br>ATPase domain of HSP90 chaperone/DNA topoisomerase II/histidine kinase<br>OS06G0622000 PROTEIN |
| evm.model.Hap1-Super-Scaffold_100036.43 | 0.5 | X | X | V | V | X | 951 | LeuD/IlvD-like<br>ACONITASE/IRON-RESPONSIVE ELEMENT FAMILY MEMBER<br>aconitase_1: aconitate hydratase 1<br>Aconitase family signature 1.<br>Aconitase C-terminal domain<br>Aconitase family (aconitate hydratase)<br>Aconitase iron-sulfur domain<br>Aconitase family signature 2. |
| evm.model.Hap1-Super-Scaffold_100036.94.1.61231383 | 0.492551118337242 | Out CMD2 contig | Out CMD2 contig | V | V | X | 414 | Sodium/calcium exchanger protein<br>caca2: calcium/proton exchanger<br>cax: calcium/proton exchanger<br>VACUOLAR CATION/PROTON EXCHANGER<br>VACUOLAR CALCIUM ION TRANSPORTER |
| evm.model.Hap1-Super-Scaffold_100036.75 | 0.4908418454937844 | Out CMD2 contig | V | V | V | X | 539 | BNAC03G34680D PROTEIN<br>NAD(P)-binding Rossmann-fold domains<br>Aminoacid dehydrogenase-like- N-terminal domain<br>aroE: shikimate dehydrogenase<br>Shikimate dehydrogenase substrate binding domain<br>Shikimate 5'-dehydrogenase C-terminal domain<br>Leucine Dehydrogenase<br>SHIKIMATE DEHYDROGENASE<br>Shikimate / quinate 5-dehydrogenase<br>DHQase_I<br>Aldolase class IAldolase<br>aroD: 3-dehydroquinate dehydratase- type I<br>Type I 3-dehydroquinase<br>NAD_bind_Shikimate_DH |

| | | | | | | | |
|---|---|---|---|---|---|---|---|
| evm.model.Hap1-Super-Scaffold_100036.87 | 0.47638444196707236 | V | V | V | V | X | 677 | E3 UBIQUITIN-PROTEIN LIGASE COP1<br>WD40 repeat-like<br>Zinc finger RING-type profile.<br>ring_2<br>RING-HC_COP1<br>Zinc/RING finger domain<br>Trp-Asp (WD) repeats signature.<br>RING/U-box<br>Zinc finger RING-type signature.<br>WD40<br>Trp-Asp (WD) repeats profile.<br>consensus disorder prediction<br>Coil<br>Zinc finger- C3HC4 type (RING finger)<br>WD domain- G-beta repeat |
| evm.model.Hap1-Super-Scaffold_100036.51 | 0.4723144809183478 | V | X | X | X | V | 312 | 2-ALKENAL REDUCTASE (NADP(+)-DEPENDENT)-LIKE<br>GroES-like<br>NAD(P)-binding Rossmann-fold domains<br>PROSTAGLANDIN REDUCTASE<br>PKS_ER_names_mod<br>Zinc-binding dehydrogenase |
| evm.model.Hap1-Super-Scaffold_100036.57 | 0.4717107073085255 | X | X | X | X | V | 607 | ZINC FINGER CW-TYPE COILED-COIL DOMAIN PROTEIN 3.<br>Zinc finger CW-type profile.<br>ATPase domain of HSP90 chaperone/DNA topoisomerase II/histidine kinase<br>OS06G0622000 PROTEIN<br>consensus disorder prediction |
| evm.model.Hap1-Super-Scaffold_100036.79 | 0.4441556977397326 | V | V | V | V | X | 611 | G1/S-SPECIFIC CYCLIN-E PROTEIN<br>consensus disorder prediction<br>OS05G0597400 PROTEIN |
| evm.model.Hap1-Super-Scaffold_100036.45 | 0.43873609978498446 | X | X | X | X | V | 505 | TERPENE SYNTHASE 12-RELATED<br>Terpene_cyclase_plant_C1<br>Terpene synthase- N-terminal domain<br>Terpene synthase family- metal binding domain<br>Farnesyl Diphosphate Synthase<br>Terpenoid synthases<br>OS04G0344100 PROTEIN-RELATED<br>TERPENE SYNTHASE 12-RELATED<br>Terpenoid cyclases/Protein prenyltransferases |
| evm.model.Hap2-Super-Scaffold_100004.514 | 0.4285325850314853 | X | X | X | X | V | 583 | G1/S-SPECIFIC CYCLIN-E PROTEIN<br>OS05G0597400 PROTEIN |

| | | | | | | | | |
|---|---|---|---|---|---|---|---|---|
| evm.model.Hap2-Super-Scaffold_100004.534 | 0.4245975125511343 | V | V | X | X | V | 759 | P-loop containing nucleoside triphosphate hydrolases<br>TRANSCRIPTION TERMINATION FACTOR 2-RELATED<br>DEXDc_SHPRH-like<br>ultradead3<br>P-loop containing nucleoside triphosphate hydrolases<br>consensus disorder prediction<br>Superfamilies 1 and 2 helicase ATP-binding type-1 domain profile. SNF2 family N-terminal domain |
| evm.model.Hap1-Super-Scaffold_100036.81 | 0.41764803912629644 | X | X | X | X | V | 372 | ARM REPEAT SUPERFAMILY PROTEIN<br>consensus disorder prediction |
| evm.model.Hap1-Super-Scaffold_100036.44 | 0.4174312036070787 | V | X | X | X | V | 394 | Terpene synthase family- metal binding domain<br>TERPENE SYNTHASE 12-RELATED<br>Terpenoid cyclases/Protein prenyltransferases<br>Farnesyl Diphosphate Synthase<br>OS04G0344100 PROTEIN-RELATED<br>Terpene synthase- N-terminal domain<br>Terpenoid synthases |
| evm.model.Hap2-Super-Scaffold_100004.494 | 0.4129948869252456 | V | V | X | V | V | 876 | consensus disorder prediction<br>OS06G0608100 PROTEIN |
| evm.model.Hap1-Super-Scaffold_100036.49.1.612312ec | 0.40287570704406056 | V | V | V | V | X | 862 | Ribonuclease H-like<br>35exoneu6<br>3'-5' exonuclease<br>HRDC domain<br>consensus disorder prediction<br>PROTEIN RRP6-LIKE 3<br>POLYMYOSITIS/SCLERODERMA AUTOANTIGEN-RELATED |
| evm.model.Hap2-Super-Scaffold_100004.502 | 0.39753655150895717 | Out CMD2 contig | Out CMD2 contig | X | X | V | 568 | E3 UBIQUITIN-PROTEIN LIGASE KEG-LIKE<br>Transferase(Phosphotransferase) domain 1<br>consensus disorder prediction<br>Protein kinase domain profile.<br>Protein kinase-like (PK-like)<br>OS06G0639500 PROTEIN |
| evm.model.Hap2-Super-Scaffold_100004.505 | 0.3967982847427726 | Out CMD2 contig | Out CMD2 contig | V | V | X | 367 | BNAA08G30680D PROTEIN<br>Triose-phosphate Transporter family<br>SOLUTE CARRIER FAMILY 35 |
| evm.model.Hap2-Super-Scaffold_100004.512 | 0.36334890110022694 | V | V | V | V | X | 580 | ARM repeat<br>consensus disorder prediction<br>ARM REPEAT SUPERFAMILY PROTEIN |

| | | | | | | | | |
|---|---|---|---|---|---|---|---|---|
| evm.model.Hap1-Super-Scaffold_100036.52.1.61231304 | 0.33391892158072123 | V | V | X | V | V | 252 | Prephenate dehydratase signature 2.<br>Periplasmic binding protein-like II<br>ACT_CM-PDT<br>ACT domain profile.<br>PREPHENATE DEHYDRATASE P PROTEIN<br>Prephenate dehydratase domain profile.<br>AROGENATE/PREPHENATE DEHYDRATASE<br>PBP2_Ct-PDT_like |
| evm.model.Hap2-Super-Scaffold_100004.555 | 0.30197426014393786 | Out CMD2 contig | Out CMD2 contig | V | V | X | 710 | HOX_1<br>Bet v1-like<br>'Homeobox' domain signature.<br>START_ArGLABRA2_like<br>Bet v1-like<br>START domain profile.<br>HOMEOBOX-LEUCINE ZIPPER PROTEIN MERISTEM L1<br>consensus disorder prediction<br>START_1<br>Coil |
| evm.model.Hap1-Super-Scaffold_100036.84 | 0.2985079020042521 | V | V | V | V | X | 194 | OXOGLUTARATE/IRON-DEPENDENT DIOXYGENASE<br>1-AMINOCYCLOPROPANE-1-CARBOXYLATE OXIDASE<br>Clavaminate synthase-like Coil |
| evm.model.Hap2-Super-Scaffold_100004.562 | 0.27981425800892473 | V | V | X | X | V | 329 | secretory_peroxidase<br>Plant peroxidase signature<br>Haem peroxidase superfamily signature<br>Peroxidases proximal heme-ligand signature.<br>Plant heme peroxidase family profile.<br>PEROXIDASE 25-RELATED<br>Peroxidases active site signature.<br>Heme-dependent peroxidases |
| evm.model.Hap1-Super-Scaffold_100036.100 | 0.2759093145551588 | V | V | V | V | X | 529 | OS07G0633600 PROTEIN |
| evm.model.Hap1-Super-Scaffold_100036.70 | 0.2625726676456461 | X | X | X | X | V | 270 | Protein tyrosine and serine/threonine kinase<br>Phosphorylase Kinase- domain 1<br>CHITIN ELICITOR RECEPTOR KINASE 1-RELATED<br>Transferase(Phosphotransferase) domain 1<br>Protein kinase-like (PK-like)<br>ILK<br>Serine/Threonine protein kinases active-site signature.<br>Protein kinase domain profile.<br>serkin_6 |

| | | | | | | | | |
|---|---|---|---|---|---|---|---|---|
| evm.model.Hap2-Super-Scaffold_100004.540 | 0.2578879414476308 | Out CMD2 contig | Out CMD2 contig | V | V | X | 395 | Galactosyltransferase<br>BETA-1-3-N-ACETYLGLUCOSAMINYLTRANSFERASE<br>Domain of unknown function (DUF4094)<br>HEXOSYLTRANSFERASE<br>Coil |
| evm.model.Hap1-Super-Scaffold_100036.59 | 0.24867590395999326 | X | X | X | X | V | 208 | Protein kinase domain profile.<br>MAP KINASE KINASE KINASE<br>Transferase(Phosphotransferase) domain 1<br>CCR4-NOT TRANSCRIPTIONAL COMPLEX SUBUNIT CAF120-RELATED<br>serkin_6<br>Protein kinase-like (PK-like)<br>Transferase(Phosphotransferase) domain 1 |
| evm.model.Hap2-Super-Scaffold_100004.490 | 0.24129512153502283 | V | V | V | V | X | 542 | OS07G0633600 PROTEIN |
| evm.model.Hap2-Super-Scaffold_100004.530 | 0.23547113417844592 | X | X | X | X | V | 392 | Protein kinase domain<br>Protein kinases ATP-binding region signature.<br>Transferase(Phosphotransferase) domain 1<br>Protein kinase domain profile.<br>MITOGEN-ACTIVATED PROTEIN KINASE KINASE KINASE 15<br>Protein kinase-like (PK-like)<br>serkin_6<br>STKc_MAPKKK<br>CCR4-NOT TRANSCRIPTIONAL COMPLEX SUBUNIT CAF120-RELATED |
| evm.model.Hap2-Super-Scaffold_100004.524 | 0.21552478682001472 | X | X | X | X | V | 577 | Transferase(Phosphotransferase) domain 1<br>CHITIN ELICITOR RECEPTOR KINASE 1<br>LysM domain profile.<br>Protein kinases ATP-binding region signature.<br>Protein tyrosine and serine/threonine kinase<br>serkin_6<br>TonB-dependent receptor proteins signature 1.<br>LysM<br>Phosphorylase Kinase- domain 1<br>CHITIN ELICITOR RECEPTOR KINASE 1-RELATED<br>Serine/Threonine protein kinases active-site signature.<br>Protein kinase-like (PK-like)<br>LysM_2<br>Protein kinase domain profile. |

| | | | | | | | | |
|---|---|---|---|---|---|---|---|---|
| evm.model.Hap1-Super-Scaffold_100036.104.1.612313d1 | 0.2112464684253818 | V | V | V | V | X | 284 | Coil<br>IGPS<br>Ribulose-phoshate binding barrel<br>TRYPTOPHAN BIOSYNTHESIS PROTEIN<br>ALDOLASE-TYPE TIM BARREL FAMILY PROTEIN-RELATED<br>Aldolase class I<br>Indole-3-glycerol phosphate synthase signature. |
| evm.model.Hap1-Super-Scaffold_100036.64 | 0.20910290552304106 | V | V | X | X | V | 427 | Phosphorylase Kinase- domain 1<br>Protein kinases ATP-binding region signature.<br>serkin_6<br>Protein kinase-like (PK-like)<br>CCR4-NOT TRANSCRIPTIONAL COMPLEX SUBUNIT CAF120-RELATED<br>Transferase(Phosphotransferase) domain 1<br>Protein kinase domain<br>MITOGEN-ACTIVATED PROTEIN KINASE KINASE KINASE 15<br>STKc_MAPKKK |
| evm.model.Hap1-Super-Scaffold_100036.46 | 0.19561513682528847 | V | V | V | V | X | 276 | Activating enzymes of the ubiquitin-like proteins<br>ThiF family<br>UBIQUITIN-ACTIVATING ENZYME E1<br>NEDD8-ACTIVATING ENZYME E1 REGULATORY SUBUNIT |
| evm.model.Hap1-Super-Scaffold_100036.33 | 0.18864454807486822 | V | V | V | V | X | 647 | HOX_1<br>Bet v1-like<br>Homeodomain-like<br>START_ArGLABRA2_like<br>Bet v1-like<br>'Homeobox' domain profile.<br>START domain profile.<br>consensus disorder prediction<br>HOMEOBOX-LEUCINE ZIPPER PROTEIN MERISTEM L1<br>START domain<br>Coil |
| evm.model.Hap1-Super-Scaffold_100036.37 | 0.18542350860571996 | X | X | X | X | V | 632 | Ligand-gated ion channel<br>Receptor family ligand binding region<br>IONOTROPIC GLUTAMATE RECEPTOR<br>Voltage-gated potassium channels<br>GluR_14<br>GLUTAMATE RECEPTOR 3.7<br>Periplasmic binding protein-like II |

| | | | | | | | | |
|---|---|---|---|---|---|---|---|---|
| evm.model.Hap2-Super-Scaffold_100004.519 | 0.17898179872411138 | V | X | V | X | X | 500 | Type I 3-dehydroquinase<br>Aldolase<br>Shikimate dehydrogenase substrate binding domain<br>Aminoacid dehydrogenase-like- N-terminal domain<br>NAD_bind_Shikimate_DH<br>NAD(P)-binding Rossmann-fold domains<br>DHQase_I<br>Shikimate 5'-dehydrogenase C-terminal domain<br>Aldolase class I<br>BNAC03G34680D PROTEIN<br>Leucine Dehydrogenase |
| evm.model.Hap1-Super-Scaffold_100036.50 | 0.1768013327966887 | V | V | X | V | V | 403 | BETA-1-3-N-ACETYLGLUCOSAMINYLTRANSFERASE<br>Coil<br>HEXOSYLTRANSFERASE<br>Galactosyltransferase |
| evm.model.Hap2-Super-Scaffold_100004.517 | 0.172929061679052 | V | V | X | X | V | 234 | ACYL-MALONYL CONDENSING ENZYME-RELATED<br>SOLUTE CARRIER FAMILY 35 MEMBER G1<br>Multidrug resistance efflux transporter EmrE |
| evm.model.Hap1-Super-Scaffold_100036.72 | 0.17067961363051942 | V | X | V | V | X | 177 | Zinc/RING finger domain<br>OS08G0421900 PROTEIN<br>FYVE/PHD zinc finger |
| evm.model.Hap1-Super-Scaffold_100036.101 | 0.1693169882843195 | V | V | V | V | X | 367 | Armadillo/plakoglobin ARM repeat profile.<br>ARM REPEAT SUPERFAMILY PROTEIN<br>U BOX DOMAIN-CONTAINING<br>Kinesin-associated protein (KAP)<br>arm_5 |
| evm.model.Hap2-Super-Scaffold_100004.520 | 0.16723068248083853 | X | X | X | X | V | 504 | ENDO-1-4-BETA-GLUCANASE<br>ENDOGLUCANASE 11<br>Glycosyl hydrolase family 9<br>Six-hairpin glycosidases |
| evm.model.Hap1-Super-Scaffold_100036.77 | 0.1633679039256505 | V | V | X | X | V | 520 | Permease family<br>XANTHINE-URACIL / VITAMIN C PERMEASE FAMILY MEMBER<br>NUCLEOBASE-ASCORBATE TRANSPORTER 2 |
| evm.model.Hap1-Super-Scaffold_100036.61 | 0.14990874525279485 | V | V | V | X | X | 222 | Coil<br>BCR-ASSOCIATED PROTEIN- BAP<br>B-CELL RECEPTOR-ASSOCIATED 31-LIKE PROTEIN-RELATED |

| | | | | | | | | |
|---|---|---|---|---|---|---|---|---|
| evm.model.Hap1-Super-Scaffold_100036.92 | 0.14846021552347244 | V | X | V | V | X | 376 | Glutaredoxin<br>Thioredoxin-like<br>PROTEIN DISULFIDE-ISOMERASE 2-3<br>pdi_dom: protein disulfide-isomerase domain<br>Thioredoxin family active site.<br>PDI_a_P5<br>P5_C |
| evm.model.Hap1-Super-Scaffold_100036.76 | 0.14564608057056844 | V | V | X | V | V | 398 | EamA-like transporter family<br>ACYL-MALONYL CONDENSING ENZYME-RELATED<br>Multidrug resistance efflux transporter EmrE<br>SOLUTE CARRIER FAMILY 35 MEMBER G1 |
| evm.model.Hap2-Super-Scaffold_100004.558 | 0.14079457843964588 | V | V | X | X | V | 186 | LATE EMBRYOGENESIS ABUNDANT (LEA) HYDROXYPROLINE-RICH GLYCOPROTEIN FAMILY |
| evm.model.Hap2-Super-Scaffold_100004.523 | 0.13604313969600076 | Out CMD2 contig | V | V | V | X | 189 | FYVE/PHD zinc finger<br>OS08G0421900 PROTEIN<br>BAH domain<br>Zinc/RING finger domain<br>CHROMATIN REMODELING PROTEIN EBS-LIKE<br>BAH domain profile.<br>BAH_4 |
| evm.model.Hap1-Super-Scaffold_100036.67 | 0.1348187373584988 | X | X | V | V | X | 484 | Cytochrome P450<br>CYTOCHROME P450 26<br>INACTIVE LINOLENATE HYDROPEROXIDE LYASE-RELATED<br>consensus disorder prediction<br>Cytochrome P450<br>E-class P450 group IV signature |
| evm.model.Hap2-Super-Scaffold_100004.485 | 0.12939014313506328 | X | X | V | V | X | 351 | Coil<br>ALDOLASE-TYPE TIM BARREL FAMILY PROTEIN-RELATED<br>Aldolase class I<br>TRYPTOPHAN BIOSYNTHESIS PROTEIN<br>Indole-3-glycerol phosphate synthase<br>Ribulose-phoshate binding barrel<br>TRYPTOPHAN BIOSYNTHESIS PROTEIN<br>IGPS<br>Indole-3-glycerol phosphate synthase signature. |
| evm.model.Hap1-Super-Scaffold_100036.48 | 0.1255629338676746 | V | V | V | V | X | 173 | UBIQUITIN-ACTIVATING ENZYME E1<br>Activating enzymes of the ubiquitin-like proteins<br>NEDD8-ACTIVATING ENZYME E1 REGULATORY SUBUNIT |

| | | | | | | | |
|---|---|---|---|---|---|---|---|
| evm.model.Hap2-Super-Scaffold_100004.500 | 0.12330663871400556 | V | X | X | V | V | 523 | ADP-glucose pyrophosphorylase signature 3.<br>LbH_G1P_AT_C<br>GLUCOSE-1-PHOSPHATE ADENYLYLTRANSFERASE<br>Nucleotide-diphospho-sugar transferases<br>ADP-glucose pyrophosphorylase signature 2.<br>Spore Coat Polysaccharide Biosynthesis Protein SpsA- Chain A<br>glgC: glucose-1-phosphate adenylyltransferase<br>Hexapeptide repeat proteins<br>Trimeric LpxA-like enzymes<br>ADP_Glucose_PP<br>Nucleotidyl transferase<br>ADP-glucose pyrophosphorylase signature 1.<br>GLUCOSE-1-PHOSPHATE ADENYLYLTRANSFERASE-RELATED |
| evm.model.Hap1-Super-Scaffold_100036.74 | 0.12014386492632803 | V | V | V | V | X | 484 | ENDO-1-4-BETA-GLUCANASE<br>Glycosyl hydrolases family 9 (GH9) active site signature 2.<br>ENDOGLUCANASE 11<br>Six-hairpin glycosidases |
| evm.model.Hap1-Super-Scaffold_100036.56 | 0.11914843783224532 | X | X | X | X | V | 182 | BCR-ASSOCIATED PROTEIN- BAP<br>Coil<br>ENDOPLASMIC RETICULUM TRANSMEMBRANE PROTEIN 3 |
| evm.model.Hap1-Super-Scaffold_100036.39 | 0.11741771868679705 | V | V | X | X | V | 191 | Vaccinia Virus protein VP39<br>SAM-dependent O-methyltransferase class II-type profile.<br>O-METHYLTRANSFERASE<br>CAFFEIC ACID 3-O-METHYLTRANSFERASE 1-LIKE<br>S-adenosyl-L-methionine-dependent methyltransferases |
| evm.model.Hap2-Super-Scaffold_100004.544 | 0.11637720642979217 | V | V | V | V | X | 99 | Activating enzymes of the ubiquitin-like proteins<br>NEDD8-ACTIVATING ENZYME E1 REGULATORY SUBUNIT<br>ThiF family<br>UBIQUITIN-ACTIVATING ENZYME E1 |
| evm.model.Hap1-Super-Scaffold_100036.105 | 0.11413109571296812 | V | V | X | V | V | 413 | SAWADEE PROTEIN |

| | | | | | | | | |
|---|---|---|---|---|---|---|---|---|
| evm.model.Hap1-Super-Scaffold_100036.95 | 0.11213985175161964 | V | V | V | V | X | 523 | ADP-glucose pyrophosphorylase signature 3.<br>LbH_G1P_AT_C<br>Hexapeptide repeat proteins<br>GLUCOSE-1-PHOSPHATE ADENYLYLTRANSFERASE<br>Nucleotide-diphospho-sugar transferases<br>ADP-glucose pyrophosphorylase signature 2.<br>Spore Coat Polysaccharide Biosynthesis Protein SpsA- Chain A<br>glgC: glucose-1-phosphate adenylyltransferase<br>Trimeric LpxA-like enzymes<br>consensus disorder prediction<br>Nucleotidyl transferase<br>ADP-glucose pyrophosphorylase signature 1. |
| evm.model.Hap1-Super-Scaffold_100036.88 | 0.11118942366578712 | X | X | X | X | V | 408 | LYSOSOMAL ACID LIPASE-RELATED LIPASE<br>Partial alpha/beta-hydrolase lipase region<br>Steryl_ester_lip<br>alpha/beta-Hydrolases<br>Serine aminopeptidase- S33 |
| evm.model.Hap1-Super-Scaffold_100036.29 | 0.10791899309684382 | X | X | X | X | V | 255 | Class II aaRS ABD-related<br>RIBOSOME BIOGENESIS PROTEIN BRX1 HOMOLOG 2<br>consensus disorder prediction<br>Brix domain profile. |
| evm.model.Hap2-Super-Scaffold_100004.491 | 0.10704299140096113 | X | X | V | V | X | 468 | RIBONUCLEASE E/G-LIKE PROTEIN-CHLOROPLASTIC<br>Ribonuclease E/G family<br>RIBONUCLEASE |
| evm.model.Hap2-Super-Scaffold_100004.510 | 0.10371731560815957 | V | V | X | X | V | 406 | Partial alpha/beta-hydrolase lipase region<br>LIPASE<br>alpha/beta-Hydrolases<br>LYSOSOMAL ACID LIPASE-RELATED<br>Steryl_ester_lip<br>Serine aminopeptidase- S33 |
| evm.model.Hap2-Super-Scaffold_100004.511 | 0.10339571768341126 | X | X | X | X | V | 272 | LYSOSOMAL ACID LIPASE-RELATED<br>alpha/beta hydrolase fold<br>alpha/beta-Hydrolases<br>LIPASE |

| | | | | | | | | |
|---|---|---|---|---|---|---|---|---|
| evm.model.Hap2-Super-Scaffold_100004.548 | 0.10282390227875222 | V | Out CMD2 contig | X | X | V | 304 | SAM-dependent O-methyltransferase class II-type profile.<br>O-methyltransferase domain<br>Dimerisation domain<br>CAFFEIC ACID 3-O-METHYLTRANSFERASE 1-LIKE<br>Vaccinia Virus protein VP39<br>""winged helix"" repressor DNA binding domain<br>O-mtase<br>O-METHYLTRANSFERASE<br>"Winged helix" DNA-binding domain<br>S-adenosyl-L-methionine-dependent methyltransferases |

**Supplemental Table 4: 81 resistance proteins of the *CMD2* region.** The resistance proteins were inferred from the CMD2 region of the 2 haplotypes of TME3. The probability of resistance was computed with prPred. Presence in 60444 genome (GCA_963409065.1 Cornet) and in previous TME3 genome (GCA_003957995.1 Kuon) (and expression (IsoSeq) were obtained after orthologous enrichment and manual analyses of phylogenetic trees. Resistance patterns of Interproscan are reported in the last column.

| Gene ID | prped probability | 60444 presence | NCBI TME3 presence | 60444 Isoseq | TME3 Isoseq | Gene Length | Function | Reported resistance |
|---|---|---|---|---|---|---|---|---|
| evm.model.Hap1-Super-Scaffold_100036.36 | 0.86 | V | X | X | V | 896 | Glumamate receptor 3.4 | Resistance to necrotrophic pathogens (Zhu et al., 2021) (74) |
| evm.model.Hap1-Super-Scaffold_100036.73 | 0.65 | V | X | X | V | 562 | NA | NA |
| evm.model.Hap2-Super-Scaffold_100004.532 | 0.53 | V | V | X | V | 613 | MORC type protein | Resistance to Turnip Crinkle Virus (TCV) in Arabidopsis, as a component of hypersensitive response to TCV (HRT) (Koch et al., 2017) (75) |
| evm.model.Hap2-Super-Scaffold_100004.494 | 0.41 | V | V | X | V | 876 | Cytochrome P450 | Xenobiotics detoxification (Pandian et al., 2020) (76) |
| evm.model.Hap1-Super-Scaffold_100036.52.1.61231304 | 0.34 | V | V | X | V | 252 | Arogenate dehydratase | Pathway of Phenylalanine biosynthesis, involved in sugarcane mosaic virus resistance |

| | | | | | | | | |
|---|---|---|---|---|---|---|---|---|
| | | | | | | | | (Yuan et al., 2019) (77) |
| evm.model.Hap1-Super-Scaffold_100036.50 | 0.18 | V | V | X | V | 403 | Hexosyltransferase | Downregulation involved in reduction to Tobacco mosaic virus resistance (Chong et al., 2022) (78) |
| evm.model.Hap1-Super-Scaffold_100036.76 | 0.15 | V | V | X | V | 398 | NA | NA |
| evm.model.Hap2-Super-Scaffold_100004.500 | 0.12 | V | X | X | V | 523 | NA | NA |
| evm.model.Hap1-Super-Scaffold_100036.105 | 0.11 | V | V | X | V | 413 | NA | NA |

**Supplemental Table 5: Putative resistance proteins of TME3.** The resistance proteins were inferred from the *CMD2* of the 2 haplotypes of TME3. The probability of resistance was computed with prPred. Presence (in genomes) and expression (IsoSeq) were obtained after orthologous enrichment and manual analyses of phylogenetic trees. Resistance patterns of Interproscan are available in Supplemental Table 4.

**Supplemental Table 6: Overview of CMD resistance markers used in this study.**

| Marker | Sequence | Type | Study |
|---|---|---|---|
| SSRY 28 Fw | TTGACATGAGTGATATTTTCTTGAG | SSR | Akano et al., 2002 |
| SSRY 28 Rev | GCTGCGTGCAAAACTAAAAT | | |
| SSRY 106 Fw | GGAAACTGCTTGCACAAAGA | SSR | Lokko et al., 2005 |
| SSRY 106 Rev | CAGCAAGACCATCACCAGTTT | | |
| SSR NS158 Fw | GTGCGAAATGGAAATCAATG | SSR | |

| | | | |
|---|---|---|---|
| SSR NS158 Rev | TGAAATAGTGATACATGCAAAAGGA | | Okogbenin et al., 2007 |
| SSR NS169 Fw | GTGCGAAATGGAAATCAATG | | |
| SSR NS169 Rev | GCCTTCTCAGCATATGGAGC | | |
| RFLP RME -1 Fw | ATGTTAATGTAATGAAAGAGC | RFLP | |
| RFLP RME-1 Rev | AGAAGAGGGTAGGAGTTATGT | | |
| SSR_NS198 Fw | TGCAGCATATCAGGCATTTC | SSR | Okogbenin et al., 2012 |
| SSR_NS198 Rev | TGGAAGCATGCATCAAATGT | | |
| s05214_1427095_chromosomeXII_5549883_23.57_- | GCTTCAAAACACTCCAGACGCTGCACAAACGTCTCTGCTGCGAAATCCCCAGTTGAAGAATGGAGGGATTGGTTTCAACTAGGGTTTCCCTCAGGACTACTC | SNP | Rabbi et al., 2014 |
| s05214_1380239_chromosomeXII_5596739_22_- | TTCCCTTTTGCCCCATGTTAAGATGCCTCCCATTTATCTAAGGCCCAATTCTGGAATTAAGGCCAAGTTGTTTTTGTTAGAGTTGTTGTAAAAGGCTGCATT | | |
| s05214_1373138_chromosomeXII_5603840_23.92_- | AAAGAAAACTGCACATGCCTACCTCAACTAACTTAAAATGGTCAATATCTGGTTTACAGACGAATGCCCCAAATACCCACTGGCAAAATCTTCTCCAGACTCC | | |
| s05214_1273143_chromosomeXII_5703835_23.57_- | GCTGCATAACTGAGGATGATCCAGCTGGGTGTTTATTTGATAGTTGATATCACCAGCTGTATGCATGATGATTTTGGTTATTCAATCCAGTAAATTGACTCC | | |
| s05214_1215853_chromosomeXII_5761125_27.55_- | TTTATTAGCTCAGTTGCATCCACTCCGATTCCTTTCACCTCCATTTGATCAGTCCTTTGCAAGCGCAACAGATGCCACCCCTTAATCATCACATGTATATAC | | |

| | |
|---|---|
| s05214_1190358_chromosomeXII_5786620_27.33_- | GAAAAATAAATGATAAGAAGAAAAGGGTATGATTCAATATCCTCATCTCTTGGCTGCTTACTCTGTTTCTGCTTCTCCTCTCTTTATACAATAATGGAATAG |
| s05214_1115740_chromosomeXII_5861238_27.33_- | TTGATTTTCTTTTTTCTTTTTTCTTTTTTCTTTTTTGCCTCCATTCCTACCTGCTACGGGGTGTAATAGCTGCGCGTAGAAGTTAAATGAGAGGTAAAATTA |
| s05214_1115005_chromosomeXII_5861973_27.33_- | ATCTTCTAATGATGCCATCTGTTGCAGCAAAACCCAATTCATCAAATTCCGAATACCCACAAATATAACACATTGAAATTATATGCATACGTATACAATCCC |
| s05214_1081665_chromosomeXII_5895313_27.33_- | TAATCTCTGCCACCAGGCATAATTCACACATCTTTTCAAGAAGCCACTAAGTTCACAATGAACAAATTGCTGCATGTAAAACTTAAATGGTAACCAATGGAC |
| s05214_1081652_chromosomeXII_5895326_27.33_- | AGTTCTACAATATTAATCTCTGCCACCAGGCATAATTCACACATCTTTTCAGAGAAGCCACTAAGTCACAATGAACAAATTGCTGCATGTAAAACTTAAATG |
| s05214_1081274_chromosomeXII_5895704_27.33_- | CATACAAACCAAGGTAACGTAAATGCAGCACCTATGGAGATGGATCTGCAGATGATTGGCAACTGAAGCCTATTAGCAAGGCACCTAACCTGCTGGTCTTTA |
| s05214_1076643_chromosomeXII_5900335_27.33_- | ACCATTTTGGTCCACTTCATCCACATGGGAATGGATACGGATGAAACCAATAGTATCTGATAACTGGGCGAGCACCAGTCAAGTTCCTTTTCCCTGTTGAAT |
| s05214_1076642_chromosomeXII_5900336_27.33_- | AACCATTTTGGTCCACTTCATCCACATGGGAATGGATACGGATGAAACCAAGTGTATCTGATAACTGGGCGAGCACCAGTCAAGTTCCTTTTCCCTGTTGAA |
| s05214_1076455_chromosomeXII_5900523_27.33_- | TCTGGTATTCAACAAAGTGACAGATTGACTAGTTCCCACCTTACTAAATCAGAGTAAAAAGAAAGTAAACAAGTAGCACACCAAGTGATACTACCTCAGGCA |

| | |
|---|---|
| s05214_1076425_chromosomeXII_5900553_28.64_- | GCAGCCAAGGAGCCAAAATAGCTTAAATTTTCTGGTATTCAACAAAGTGACCAGAGATTGACTAGTTCCCACCTTACTAAATCAAGTAAAAAGAAAGTAAACAA |
| s05214_1041782_chromosomeXII_5935196_28.82_- | CTAATTAACATCTCTCACAGAATCCATGTGATAGTTCTTTCGTTCTTGCTTCCTTTATATGCACATATGCAGACAATCAAGTTCCAGGCAGAAACGGGAAGC |
| s05214_1011668_chromosomeXII_5965310_28.2_- | CTAGACAGGATGTGTCTTTGTTATCACCAAATAATACTGATAGTCTCTGCAGATTCCATGGTGGAGACACAAGGATATATTTGCCAAGAACATAAAGTACCG |
| s05214_981075_chromosomeXII_5995903_33.36_- | GGGCCATAGTTCCTGCGGTGAAGCCGGAGCCTAGACCATCTCCTTCCTTCGACTGCGGCAGCAGCGGCTTCTCCGGATCAAACCGCGCAGGCGGGGGCGAAG |
| s05214_980760_chromosomeXII_5996218_28.64_- | TGTTTTGGCCTTTGGCAGCGGATTATAAAACATAAATCACTGGTTTTTTCGGTTTGTTTTTTTTTTTAAAACATAAATCACCGGTAACCATGGGCAAATGAACAAA |
| s05214_980758_chromosomeXII_5996220_26.75_- | TCTGTTTTGGCCTTTGGCAGCGGATTATAAAACATAAATCACTGGTTTTTTCGTTTTTTTTTTTTAAAACATAAATCACCGGTAACCATGGGCAAATGAACAAA |
| s05214_957480_chromosomeXII_6019498_29.27_- | ATAGGAAATAGATATTGATAGTGATGCTATGGACTGAAAGGGGGAATTGGCGTCAGCAGCATCTTAAGCGTAGTGACATAGTTGGCATATTTTTAACTAGTC |
| s05214_864519_chromosomeXII_6112459_38.83_- | TAATCTATTCTTTATTTTTTGTAATTATGGAAGTGGAAGTTTTGTAAAGAGAAGAGAGGCCGTCTTCCCATCTTCAAAATTCAAAAGAAAATAAATAAAATTAC |
| s05214_776142_chromosomeXII_6200836_38.13_- | TGTGATGTTTTTGAAGTTTTTGAGAGAGCGAGCTCGGATTATTACTACTTAGTTGTGCTCTAGCTTTGGCTGAATGCTATTTGTTTTCAGTTTCCCAGTTTT |

| | |
|---|---|
| s05214_755846_chromosomeXII_6221132_29.27_- | CTTGATGGTCTTTTTGGAACAGGAACTGGTTCCACTGTTTCTGTGTAAGTTATAAACTTTTTTGAGCCGTCTTTAGCAGCTTCTTTTATACAGAAATGTCGAT |
| s05214_755621_chromosomeXII_6221357_37.11_- | GTATTAAGCAGAGGGATTGGCTGGCAGGTGTGCTGGTTTAAAAGTAATTGCTTTTTGGTTGAGCATGTAGGAAATAATATTACAATCCAGTGCAAAATTTTT |
| s05214_730604_chromosomeXII_6246374_29.27_- | AAAATTTGGTATATTTGGCCCTCGTTGTATAGTTCAGGGGTAAATTGGCCTCTTTAGCCGTTACCAAACTACCTGTCTAAGAATTGCAGCCCTCCTAGTAAA |
| s05214_730219_chromosomeXII_6246759_38.13_- | TTGAGGTACTATTTTGTTCTGCCTTTCCCTATTCTCTCTCTCGGCATGCAAGATTCCATTAGGATCTGGAACTATGTTCCCATTTATGTTCTTTAGATCATAA |
| s05214_730196_chromosomeXII_6246782_38.13_- | CAGAGCTGCAAATGTCCAGTATATTGAGGTACTATTTTGTTCTGCCTTTCCGCTATTCTCTCTCTCGGCATGCAAGTTCCATTAGGATCTGGAACTATGTTC |
| s05214_719298_chromosomeXII_6257680_30.25_- | CAGAATACTCATTTTTTCCCTGTGAAGATGTGTTGCATCTTGACAGGATCCACATGATCATAGATTCATCTGCAATCTTGCAATTCTGCATTAAGCAATGCA |
| s05214_719029_chromosomeXII_6257949_30.25_- | ATTATTTGGAGCAGCAAGTGCTAGGAAACTTCTGTTTTGGAGAGCCTTAACTGGGATATCTCTCATTGTTGAGTTTTATTTATTGGTAAGTTCAACTTGGTG |
| s05214_656337_chromosomeXII_6320641_37.19_- | CCCAATGTCTGCTAACTCTAAAGGCCAAACTAAAAATTTCTCTATTTCTTCCTTCTTTTCTTCTCCAAACCTCATACATGGATGGGATAGCGACTCCATAGTG |
| s05214_656334_chromosomeXII_6320644_37.19_- | TTTCCCAATGTCTGCTAACTCTAAAGGCCAAACTAAAAATTTCTCTATTTCCTTTTCTCTTTTCTTCTCCAAACCTCATACATGGATGGGATAGCGACTCCATA |

| | |
|---|---|
| s05214_656308_chromosomeXII_6320670_36.65_- | ACAGACTTTAGGGTCTAAAAGCAGCGTTTCCCAATGTCTGCTAACTCTAAAGGGCCAAACTAAAAATTTCTCTATTTCTTCTCTTTTCTTCTCCAAACCTCA |
| s05214_611967_chromosomeXII_6365011_42.72_- | TTCTTCTTGTTCTCATAAGTTGAATCACCTAGTGTAGCTGAGGCGCGCCTCTGGGGGCACCTTGCGCCTGGGTGCGCCTTGGCTGCCTCGTCTCGCCTCCAG |
| s05214_472417_chromosomeXII_6504561_38.13_- | TTGTAAACATTTATAATCTGCAAGTTTAAAACGTAAATTACTTGAGCAAGACATTTCCAATTAAAAAAGAAAATCAACCATGCATGCAATGTTCTACCTTTAA |
| s05214_472404_chromosomeXII_6504574_38.13_- | TTCGTACTCCTGATTGTAAACATTTATAATCTGCAAGTTTAAAACGTAAATGTACTTGAGCAAGACTTTCCAATTAAAAAAGAAAATCAACCATGCATGCAA |
| s05214_472398_chromosomeXII_6504580_38.13_- | AGCGCTTTCGTACTCCTGATTGTAAACATTTATAATCTGCAAGTTTAAAACGGTAAATTACTTGAGCAAGACTTTCCAATTAAAAAAGAAAATCAACCATGC |
| s05214_371707_chromosomeXII_6605271_39.18_- | TACCTAGCTGAAACAGAACTTCCTGACCCCATAATATTCGCTTCTGCAGAGACCATTGTGATTGTGAAAAGTTCCTGAAAAATAATCCTGCTGTGAAAAGCC |
| s05214_371683_chromosomeXII_6605295_39.18_- | TAAGCAAGGCAGCCTATGTATACATACCTAGCTGAAACAGAACTTCCTGACTCCCATAATATTCGCTTCTGCAGAGCCATTGTGATTGTGAAAAGTTCCTGA |
| s05214_283847_chromosomeXII_6693131_43.86_- | GCAAGTCCAAAAAGCCTAATATCCAAGTAAATATTCACAACTTCATACCCGAAGAAAACTATGCTATGATACAACCTTCTTACAAGACCTGGAAAGGCCTTT |
| s05214_232019_chromosomeXII_6744959_38.13_- | TGGCTCTCTTGGACGACGGGGTAACCGAACTCTGATTGGACAATGAACAACTTACATTCCAAACTAGCTGCCATTGAAGAACTCCTTACCACTATTGCTAAA |

| | |
|---|---|
| s05214_232013_chromosomeXII_6744965_38.13_- | AGAATTTGGCTCTCTTGGACGACGGGGTAACCGAACTCTGATTGGACAATGTAACAACTACATTCCAAACTAGCTGCCATTGAAGAACTCCTTACCACTATT |
| s05214_232000_chromosomeXII_6744978_38.13_- | AAAATAGTGGAGAAGAATTTGGCTCTCTTGGACGACGGGGTAACCGAACTCTTGATTGGACAATGAACAACTACATTCCAAACTAGCTGCCATTGAAGAACT |
| s05214_231473_chromosomeXII_6745505_42.43_- | GATTAAAAGCCCAGCCCAGAGATTGGCCCTTAGACAATGGAAGGAATATTTCCAGAAGGTACAAGAAGAGAGGAGCGGATTGCATAGGAGCCGTTGGAACCA |
| s05214_42890_chromosomeXII_6934088_44.24_- | TATTGTGCGACAATCTACAAACGGTAAAATTATTCCTAACGTTAAATTTCTCGGCAGAACATAAATGACTAACCAGCAGCACTATAGAGTACAATGATTGCA |
| s05214_42884_chromosomeXII_6934094_42.43_- | CCATAATATTGTGCGACAATCTACAAACGGTAAAATTATTCCTAACGTTAACATTTCTGGCAGAACATAAATGACTAACCAGCAGCACTATAGAGTACAATG |
| s05214_42862_chromosomeXII_6934116_44.24_- | TTCCTCAGTGATCTCCCTTCCTCCATAATATTGTGCGACAATCTACAAACGAGTAAAATTATTCCTAACGTTAAATTTCTGGCAGAACATAAATGACTAACC |
| s05214_42565_chromosomeXII_6934413_44.24_- | AGTAGCTGAAACTGCTGCAACTTGTAGATTAGAACAGACACTTGAATCGGATGTCTGTTACTGCGACATAGTCTTTATCATCCTGTACGCCAGTTGGCTCTG |
| s05214_42557_chromosomeXII_6934421_44.24_- | TTATAAGCAGTAGCTGAAACTGCTGCAACTTGTAGATTAGAACAGACACTTAGAATCGGAGTCTGTTACTGCGACATAGTCTTTATCATCCTGTACGCCAGT |
| s05214_42485_chromosomeXII_6934493_44.24_- | TTCGGTGGACTTCTTTCACAATTTTCACATGTGTTTATGGCAGAAATTTCAGCTTTCATCTAAATTTGTATCCTTATAAGCAGTAGCTGAAACTGCTGCAAC |

| | |
|---|---|
| s05214_42147_chromosomeXII_6934831_44.24_- | GGATGCAGTGAAAGATCTGGATTCAGAGCAAGCTAAATCTTGTTAGAGATAGCATTGACCTTGATTTTATGAGTGTATCTTGGTCAAAGAAAGCATGATGAG |
| s05214_42144_chromosomeXII_6934834_44.24_- | AACGGATGCAGTGAAAGATCTGGATTCAGAGCAAGCTAAATCTTGTTAGAGAATACATTGACCTTGATTTTATGAGTGTATCTTGGTCAAAGAAAGCATGAT |
| s05214_42097_chromosomeXII_6934881_39.78_- | ATGTATTTTATTAGCGGCAGCAATGCATGGTACGCTCAATGTTTTACAACGAGATGCAGTGAAAGATCTGGATTCAGAGCAAGCTAAATCTTGTTAGAGATA |
| s06906_361646_chromosomeXII_7108969_45.41_- | CAAAAGAAAGAAAAACAAAGAATTATACAATTAATATTATGCCAAGGAACTATAAAAAAATGGTGATGACGGGAAAATGTACAGGATATAGTAGCAAAAGACT |
| s06906_361175_chromosomeXII_7109440_42.43_- | AATCTCTGGAGCTTCCACTTCCATACTCAGCTCCAGCCAAAATAATGGTGTCCATGTCCAGCAGTCTTGTACCTCTGAAACAGTAGAATGATCACCCAGAGT |
| s06906_314513_chromosomeXII_7156102_40.15_- | CATGATGACTTTGTCGTGTATCACAGCCAGAAAAGGAGCCAAGCATCCCCTCGTCGTGGTTCTTGATGAAGTATTTGGCAACCGGTGCTAAGCCAAAAAACC |
| s06906_282077_chromosomeXII_7188538_41.73_- | TTGTAAGAGATTTTGGAACGCTTCTTTGCCAGGGCATCACCCAATACAGATGATGCCACTTCTTCCATAATCATCATCTACAAAGACGGCAATTACCTCCCT |
| s06906_70963_chromosomeXII_7399652_52.51_- | GTCTCCGCCTCCAGCTTCTATAATCTTTTCTTATTTGTTACAGGGGTTTGGTTTTCAGATATCACTTAGCATACATAATGGGGAAGAAGCGAAAGCATAGTG |
| s06906_70962_chromosomeXII_7399653_52.51_- | CGTCTCCGCCTCCAGCTTCTATAATCTTTTCTTATTTGTTACAGGGGTTTGAGTTTCAGATATCACTTAGCATACATAATGGGGAAGAAGCGAAAGCATAGT |

| | | | |
|---|---|---|---|
| s06906_70919_chromosomeXII_7399696_51.59_- | TCGACGCAGCAAACAGGTCTGCTTCCTCCTCCTCTTTCGCCGCCGTCTCCGCCCTCCAGCTTCTATAATCTTTTCTTATTTGTTACAGGGGTTTGGTTTCAG | | |
| s06906_39739_chromosomeXII_7430876_52.51_- | TTTGGCACTTCAATTGTAGAGGTGAATACTCTGTCAAATCGGGCTATAGAAGTGGCCAGACAAATCAAGCTGCAACAAATGCCTAGTTCTAGTACTTCGGGC | | |
| s06906_39732_chromosomeXII_7430883_52.51_- | ACTTGGCTTTGGCACTTCAATTGTAGAGGTGAATACTCTGTCAAATCGGGCATATAGAATGGCCAGACAAATCAAGCTGCAACAAATGCCTAGTTCTAGTAC | | |
| s06906_39485_chromosomeXII_7431130_47.64_- | GAGAGTGGATTCAAAATGTTTCGGTTGAAAAGAAAAGCTTCTCTCAAAAGGCAAATAAAGAGATTTTAATCAAAGCATTTGCTTCGGCCATCTCAATCTACA | | |
| s06906_39459_chromosomeXII_7431156_52.51_- | AGCAGCAAGTTTTTAGCTTCCTCAAAGAGAGTGGATTCAAAATGTTTCGGTATGAAAAGAAAAGCTTCTCTCAAAAGGAAATAAAGAGATTTTAATCAAAGC | | |
| S12_7828503_0.69_Rabbi_et_al_2020 | CCTCCTTATAGTTTAAAAGTGTAGAATTTAGTGTATTGGGATCTTTCAAGTAATGGGTTGTTTACTATTAAGTTTGCATATGCTGCATTAGCTAATAGCTTGTGGGGTTTGGAAAATAGTGAATGGAAGTTTGCTTGGAGTTGAGCTGGCACTAAGAGTATTTATACCTTTTTCTAATTAGTGTAGTACGAGAAGTTGATA | SNP | Rabbi et al., 2020 |
| S12_7828514_0.27_Rabbi_et_al_2020 | TTTAAAAGTGTAGAATTTAGTGTATTGGGATCTTTCAAGTAATGGGTTGTTTACTATTAAGTTTGCATATGCTGCATTAGCTAATAGCTTGTGGGGTTTGGAAAATAGTGAATGGAAGTTTGCTTGGAGTTGAGCTGGCACTAAGAGTATTTATACCTTTTTCTAATTAGTGTAGTACGAGAAGTTGATAGCAGCGGATGG | | |

| | |
|---|---|
| S12_7842538_5.96_Rabbi_et_al_2020 | CGAGTAGCGGATTGCTCAACCTCTTCAAGTATAATTCTATTAAAAATTAAAGAAGAAACAAAAAATTGGCAGCAACCCTTGTGCCTGAATGGTGGTGTAAAATTGAATATTTATAAATTTCTATTCACAGTTAAGAGAAAAACTAAACTTTGGAAATGGACATATATATAAATTTTAGTAACACACACATAACAAAAAAAGA |
| S12_7842568_17.24_Rabbi_et_al_2020 | ATAATTCTATTAAAAATTAAAGAAGAAACAAAAATTGGCAGCAACCCTTGTGCCTGAATGGTGGTGTAAAATTGAATATTTATAAATTTCTATTCACAGTTAAGAGAAAAACTAAACTTTGGAAATGGACATATATATAAATTTTAGTAACACACACATAACAAAAAAAGAACACGGACAAACCTTCCTTCATCCTCCATA |
| S12_7842654_6.57_Rabbi_et_al_2020 | TTTCTATTCACAGTTAAGAGAAAAACTAAACTTTGGAAATGGACATATATATAAATTTTAGTAACACACACATAACAAAAAAAGAACACGGACAAACCTTCCTTCATCCTCCATATCTTTTTAACAATGCTTGAAACTGCAAGTTTTGCAGCATCTTGCATGGCAAGACCCTTCATTGACAATCCTATATATTGGAAATGG |
| S12_7842657_0.93_Rabbi_et_al_2020 | CTATTCACAGTTAAGAGAAAAACTAAACTTTGGAAATGGACATATATATAAATTTTAGTAACACACACATAACAAAAAAAGAACACGGACAAACCTTCCTTCATCCTCCATATCTTTTTAACAATGCTTGAAACTGCAAGTTTTGCAGCATCTTGCATGGCAAGACCCTTCATTGACAATCCTATATATTGGAAATGGCAA |
| S12_7842670_16.26_Rabbi_et_al_2020 | AGAGAAAAACTAAACTTTGGAAATGGACATATATATAAATTTTAGTAACACACACATAACAAAAAAAGAACACGGACAAACCTTCCTTCATCCTCCATATCTTTTTAACAATGCTTGAAACTGCAAGTTTTGCAGCATCTTGCATGGCAAGACCCTTCATTGACAATCCTATATATTGGAAATGGCAAGTGTGCTCTTTAT |
| S12_7849710_0.07_Rabbi_et_al_2020 | TTCAATGAATAAAAATTCATCTATGAATAGAGCAGCAGCGTCTGATCAGGAGAGCAGTGGTTGGACAGCTTATTTTGAAGATTTCTCTACCCATAGAGATCAAGATGATTGTTTCTCTTCTGGTTTTGGTAGCTCTTCAATGGTGTCTGATGCCGCATCTTATCCTGCATGGAAATCGTCAACTCATCATAATTATAACCA |

| | |
|---|---|
| S12_7874196_19.69_Rabbi_et_al_2020 | GGTCGGATTTCCGGGTCGTTACAATGAGTTTGGGAGACAGTAGCTGCTATGGCTTTGCCCTTGTTTCTATTCTTGAGTTCCATAAACCATTCAGGATATCTATGCACTCTAAAGCATGTATCCTTAGTATGACCAGTTGATTTGCAATGAATGCAAATCCTATCATCTTTCTTCCCAGATACTCCCTTCTTGAAGGCAATC |
| S12_7874273_15.64_Rabbi_et_al_2020 | TTCCATAAACCATTCAGGATATCTATGCACTCTAAAGCATGTATCCTTAGTATGACCAGTTGATTTGCAATGAATGCAAATCCTATCATCTTTCTTCCCAGATACTCCCTTCTTGAAGGCAATCTTGTTATAACCTATGTTACTCTTAGCAGCAAAAATATTGTTAATATCATCACCAGGAAGATTATGCACTTCTCTTTG |
| S12_7874328_4.68_Rabbi_et_al_2020 | CCAGTTGATTTGCAATGAATGCAAATCCTATCATCTTTCTTCCCAGATACTCCCTTCTTGAAGGCAATCTTGTTATAACCTATGTTACTCTTAGCAGCAAAAATATTGTTAATATCATCACCAGGAAGATTATGCACTTCTCTTTGTTTCTTAATCCTTAAAATCATTGAATATGCTTTGTTAACACTGGGCAAGGGATCT |
| S12_7874331_4.64_Rabbi_et_al_2020 | GTTGATTTGCAATGAATGCAAATCCTATCATCTTTCTTCCCAGATACTCCCTTCTTGAAGGCAATCTTGTTATAACCTATGTTACTCTTAGCAGCAAAAATATTGTTAATATCATCACCAGGAAGATTATGCACTTCTCTTTGTTTCTTAATCCTTAAAATCATTGAATATGCTTTGTTAACACTGGGCAAGGGATCTAGC |
| S12_7874349_4.55_Rabbi_et_al_2020 | CAAATCCTATCATCTTTCTTCCCAGATACTCCCTTCTTGAAGGCAATCTTGTTATAACCTATGTTACTCTTAGCAGCAAAAATATTGTTAATATCATCACCAGGAAGATTATGCACTTCTCTTTGTTTCTTAATCCTTAAAATCATTGAATATGCTTTGTTAACACTGGGCAAGGGATCTAGCAGCAAGATTTGATTCTTT |
| S12_7874475_17.03_Rabbi_et_al_2020 | TTCTTAATCCTTAAAATCATTGAATATGCTTTGTTAACACTGGGCAAGGGATCTAGCAGCAAGATTTGATTCTTTGCATTATCTTAGCTATCATTTAAATCCATTATAAATTGTATGAGCCTATCATCATTCTCCATTTCACACAAATCCTTTGCAGCACCACACTCACACATAGGCAGAGGCCCCAAGCATGGAAGTTCA |

| | |
|---|---|
| S12_7926088_36.43_Rabbi_et_al_2020 | TGCTAAAAGAAAAGGTCAAAAACTTTCAGCCAAGTTTTATGGACCTTTCGTGGTTCTTGAGCAAATCGGTTCCATGGCTTATAAGCTAGACTTACCTGCACACTCAAAGCTGCATCCTATTTTCCATGTTTCCACCCTCAAATGGTATCACAAAGGACAAGATTCTTGTACTCCAATGCTGCCACCAACTCCACCTGATGT |
| S12_7926132_111.76_Rabbi_et_al_2020 | CTTTCGTGGTTCTTGAGCAAATCGGTTCCATGGCTTATAAGCTAGACTTACCTGCACACTCAAAGCTGCATCCTATTTTCCATGTTTCCACCCTCAAATGGTATCACAAAGGACAAGATTCTTGTACTCCAATGCTGCCACCAACTCCACCTGATGTTCCTCTTCAACCTCTGGCTGTTTTAGACCAGCGTATTATGGCCA |
| S12_7926163_110.47_Rabbi_et_al_2020 | GGCTTATAAGCTAGACTTACCTGCACACTCAAAGCTGCATCCTATTTTCCATGTTTCCACCCTCAAATGGTATCACAAAGGACAAGATTCTTGTACTCCAATGCTGCCACCAACTCCACCTGATGTTCCTCTTCAACCTCTGGCTGTTTTAGACCAGCGTATTATGGCCAGACAACTTGAGATTTTAGTCCATTGGTGTGC |
| S12_7954248_100.35_Rabbi_et_al_2020 | CCATGGATTCACTTTTGCTCAAGATTTTGCTGATTCAATGGTTAAACTTGGTGATGTTGGCGTCCTCACTGGCTGCCAAGGTGAAATCAGGAAACGCTGCACTTTCGTCAATTAATTATTCCCCATATTTCTTCTTCTGCTTCTCCTAATAAAATAATGATTTGATCAATCCATTGATCGTATTGCAGCAAAGTTACATGT |
| S12_7954303_103.86_Rabbi_et_al_2020 | GTTGGCGTCCTCACTGGCTGCCAAGGTGAAATCAGGAAACGCTGCACTTTCGTCAATTAATTATTCCCCATATTTCTTCTTCTGCTTCTCCTAATAAAATAATGATTTGATCAATCCATTGATCGTATTGCAGCAAAGTTACATGTTTGTGTAAATTTCTTTTCAAATTTTTGTATTCTTTTTTTTTTTTTTTTAGGTTTT |
| S12_7961675_29.87_Rabbi_et_al_2020 | CTAGACCTTCACCTTCATGGTTTTGATGCAAAATGTCACATACCTCTGTCCACTATGTTGCTGCTAATAAGTAAAGACGATCTGAGTCGAAGTGGCGAGTTTATCCATATAGACACTGCATGGGGGTAGGAGAAAAAGGCATGATCAATTGTTTTCTTTTCACCACAG |

| | |
|---|---|
| | AATGCATAATGTGGATTCAAAACTTGAATTTGA |
| S12_7961873_11.06_Rabbi_et_al_2020 | TGACGATATAAAATCTCCTTCACAGGAAGCTGTTCCACATAAAGAAGCTAAAGAAACATCCGAAGCTTGGTAGAAAGGTTAAGTTTCCAGAATTGTTTCTAAAAATGGGAAGAAAGGTTATATGGAAGCTGGCCCGAAGTACTAGAACTAGGCATTTGTTGCAGCTTGATTTGTCTGGCCATTCTATAGCCCGATTTGACA |
| S12_7961874_11.1_Rabbi_et_al_2020 | GACGATATAAAATCTCCTTCACAGGAAGCTGTTCCACATAAAGAAGCTAAAGAAACATCCGAAGCTTGGTAGAAAGGTTAAGTTTCCAGAATTGTTTCTAAAAATGGGAAGAAAGGTTATATGGAAGCTGGCCCGAAGTACTAGAACTAGGCATTTGTTGCAGCTTGATTTGTCTGGCCATTCTATAGCCCGATTTGACAG |
| S12_7961875_11.37_Rabbi_et_al_2020 | ACGATATAAAATCTCCTTCACAGGAAGCTGTTCCACATAAAGAAGCTAAAGAAACATCCGAAGCTTGGTAGAAAGGTTAAGTTTCCAGAATTGTTTCTAAAAATGGGAAGAAAGGTTATATGGAAGCTGGCCCGAAGTACTAGAACTAGGCATTTGTTGCAGCTTGATTTGTCTGGCCATTCTATAGCCCGATTTGACAGA |
| S12_7961876_11.43_Rabbi_et_al_2020 | CGATATAAAATCTCCTTCACAGGAAGCTGTTCCACATAAAGAAGCTAAAGAAACATCCGAAGCTTGGTAGAAAGGTTAAGTTTCCAGAATTGTTTCTAAAAATGGGAAGAAAGGTTATATGGAAGCTGGCCCGAAGTACTAGAACTAGGCATTTGTTGCAGCTTGATTTGTCTGGCCATTCTATAGCCCGATTTGACAGAG |
| S12_7961878_10.97_Rabbi_et_al_2020 | ATATAAAATCTCCTTCACAGGAAGCTGTTCCACATAAAGAAGCTAAAGAAACATCCGAAGCTTGGTAGAAAGGTTAAGTTTCCAGAATTGTTTCTAAAAATGGGAAGAAAGGTTATATGGAAGCTGGCCCGAAGTACTAGAACTAGGCATTTGTTGCAGCTTGATTTGTCTGGCCATTCTATAGCCCGATTTGACAGAGTA |

| | |
|---|---|
| S12_7962234_32.95_Rabbi_et_al_2020 | CTGGTAAATGGAAGCAAGACATGGTGTAGATTGAGATGGCCGAAGCAAATGCTTTGATTAAAATCTCTTTATTTCCTTTTGAGAGAAGCTTTTCTTTTCAACCGAAACATTTTGAATCCACTCTCTTTGAGGAAGCTAAAAACTTGCTGCTTAGAACGAGGGACACTAGAGGGTAAACCTAAGAATTTGTCTTGAGTGTTA |
| S12_8038259_7.34_Rabbi_et_al_2020 | AGGAACTGATGGCAAGACTACCCTCTTGCACTTTGTCATCCAGGAGATTATCCGTTCTGAAGGTAAACGAGCTCTCCGCAGCATAAAAGCGAGCCAGAGTACTTGTAGTTTAAAGTCAGAGGATTTGGTTGAGGATACTAATCAGTCATCAGAACACTATCGTAACCTGGGTCTTAAGGTTATTTCAGGCTTAAGCAATGA |
| S12_8038319_28.2_Rabbi_et_al_2020 | AGGTAAACGAGCTCTCCGCAGCATAAAAGCGAGCCAGAGTACTTGTAGTTTAAAGTCAGAGGATTTGGTTGAGGATACTAATCAGTCATCAGAACACTATCGTAACCTGGGTCTTAAGGTTATTTCAGGCTTAAGCAATGAATTAGAAGACGTAAAAAATGCAGCAGCAGTAGATGCTGACGTCCTAACATCTACAGTTTC |
| S12_8039677_22.6_Rabbi_et_al_2020 | TTTCAGGTGGCGTGGATATCCTGGAGGCTTCATAACTCTATTGTACACTTTACACTGAGAAAAAAGATTCGCATACCTCAATCCTGCCTGTGGCAGCATACGTGCATGTCATTCTTGAAATTCCTTGATTTAAATGATGCTTAAAATTTTAGGGAGGTAATTTCATTATAGAAAGAGATTCACTTGGAGGAGTTCTTTGTT |
| S12_8039686_20.83_Rabbi_et_al_2020 | GCGTGGATATCCTGGAGGCTTCATAACTCTATTGTACACTTTACACTGAGAAAAAAGATTCGCATACCTCAATCCTGCCTGTGGCAGCATACGTGCATGTCATTCTTGAAATTCCTTGATTTAAATGATGCTTAAAATTTTAGGGAGGTAATTTCATTATAGAAAGAGATTCACTTGGAGGAGTTCTTTGTTAATTCCTAC |

| | |
|---|---|
| S12_8071420_30.62_Rabbi_et_al_2020 | AATCCTCTACTCCTCTCCTATATAATCTCTTTCCACCTACTGCCGAATTAAAACCCTCCATGGCTATTAAGCGTCTATCTGAAAATCCGCCACCTGCTGCTTCCTCTTCGGAAGAAGAGGAGGAGGAAGAAAACGACTCTGTGGAGAAAAACGACAGTGAGGATGAGCAAAAGGACGTCGGTGATGGAGACGACGATGAGG |
